# Naked mole-rat transcriptome signatures of socially-suppressed sexual maturation and links of reproduction to aging

**DOI:** 10.1101/221333

**Authors:** Martin Bens, Karol Szafranski, Susanne Holtze, Arne Sahm, Marco Groth, Hans A. Kestler, Thomas B. Hildebrandt, Matthias Platzer

## Abstract

Naked mole-rats (NMRs) are eusocially organized in colonies. Although breeders carry the additional metabolic load of reproduction, they are extremely long-lived and remain fertile throughout their lifespan. Comparative transcriptome analysis of ten organs from breeders and non-breeders of the eusocial long-lived NMR and the polygynous shorter-lived guinea pig provide comprehensive and unbiased molecular evidence that sexual maturation in NMR is socially suppressed. After transition into breeders, transcriptomes are markedly sex-specific, show pronounced feedback signaling via gonadal steroids and have similarities to reproductive phenotypes in African cichlid fish. Further, NMRs show functional enrichment of status-related expression differences associated with aging. Lipid metabolism and oxidative phosphorylation – molecular networks known to be linked to aging – were identified among most affected gene sets. Further, a transcriptome pattern associated with longevity is reinforced in NMR breeders contradicting the disposable soma theory of aging and potentially contributing to their exceptional long life- and healthspan.

## Introduction

The naked mole-rat (NMR, *Heterocephalus glaber*) has become increasingly popular as an animal model in a variety of research fields due to its unique biology. This includes an exceptionally long lifespan and resistance to cancer [1, 2]. According to The Animal Ageing and Longevity Database (AnAge) [3] the maximum recorded lifespan is 31 years, i.e. 368% of the prediction based on body mass. NMRs stay fertile throughout their long and healthy life, i.e. show an extraordinary long life- and healthspan [4]. This lifelong fertility becomes even more astonishing, considering the extreme reproductive skew in NMR colonies. Like eusocial insects, NMRs are socially organized in colonies consisting of a pair of reproducing animals (breeders, queen and pasha) and up to 300 subordinates (non-breeders, female and male workers) [5]. However, although workers are in principle capable of reproduction [6, 7], sexual maturation is suppressed through the behavior of the dominating queen [8]. Non-breeding animals of both sexes are the backbone of the social organization of the colony and take care of foraging, brood care, colony defense and digging [9].

Naturally, new NMR colonies originate from fissioning of existing colonies or formation of new ones by dispersers that leave their natal colony [10, 11]. When under laboratory conditions non-breeders are removed from the colony and paired with the opposite sex, they have the capability to ascend into breeders. This process is accompanied with physiological and behavioral changes, and results in the formation of a new colony [5, 6]. Remarkably, despite the queen’s enormous metabolic load of producing a large litter every three months and being exclusively in charge of lactation [12], data from wild and laboratory NMR colonies indicate that breeders live longer than their non-breeding counterparts [13, 14]. This contrasts the disposable soma theory of aging, which hypothesizes that energy is either allocated to body maintenance or reproduction [15].

In NMR, the reproductive suppression in female non-breeders is mediated through inhibition of gonadotropin-releasing hormone (GnRH) secretion from the hypothalamus [6]. This in turn leads to an inhibition of follicle stimulating hormone (FSH) and luteinizing hormone (LH) released by the pituitary gland and causes a block of ovulation. Also for male non-breeders, reproductive suppression is caused by inhibition of GnRH secretion, which results in lower levels of urinary testosterone and plasma LH [7]. The impact, however, is less profound compared to females as spermatogenesis is attenuated, but not entirely suppressed [16]. Nevertheless, weight of testis and number of active spermatozoa is higher in breeders [7, 17]. The role of GnRH in mediating environmental cues to allow or block reproduction is well described in a variety of species [18, 19].

The NMR can be regarded as a neotenic species and the prolonged retention of juvenile features has been linked to its longevity [20]. In comparison to mice, e.g. postnatal NMR brain maturation occurs at slower rate [12] and puberty is delayed. Female and male NMRs may reach sexual maturity at 7.5 to 12 months of age [21]. In the colony, however, the queen suppresses sexual maturation in both non-breeding males and females by aggressive social behavior [8] and can delay – independently of neoteny - the puberty of female workers for life [22]. Thus, sexual dimorphism is almost absent among non-breeding NMRs [23, 24]. Both sexes show almost no difference in morphology, including body mass, body size and even external genitalia, as well as no behavioral differences, in the sense that non-breeders participate and behave equally in all colony labors [25]. Nevertheless, these features are correlated with colony rank. Most profound differences can be observed comparing NMR queens vs. non-breeders, reflected in morphological differences, such as an elongated spine and higher body mass of queens, as well as behavioral differences, such as increased aggressiveness, copulation and genital nuzzling [25].

In this work, we characterized the transcript signature of reproductive status (breeder vs. non-breeder) in tissue samples of ten organs or their substructures (hereinafter called for simplicity “tissues”) from both sexes using RNA-seq. We contrast the NMR results with the transcript profiles of corresponding samples of guinea pig (GP, *Cavia porcellus*), a closely related, polygynous, not long-lived rodent species (AnAge: 12 years maximal longevity, 89% of the prediction based on body mass). We specifically focused our analyses on transcriptome signatures of the socially-suppressed sexual maturation in NMR as well as on differentially expressed genes (DEGs) that may contribute to the exceptional long life- and healthspan of NMR breeders.

## Results

To gain molecular insights into the fascinating combination of NMR phenotypes, in particular their eusocial reproduction, lifelong fertility, extraordinary healthspan and longevity, we aimed to collect a comprehensive set of tissue samples for male and female breeders and non-breeders of NMR and GP - six biological replicates each. Towards this, NMR non-breeders were removed from their natal colony, paired with an unrelated animal of the opposite sex from a second colony and thereby turned into breeders. Respective male and female litter siblings remained in the two colonies as non-breeder controls. Time to first litters averaged in 6.5±4.9 (±SD) months and duration of pregnancies was approximately 70 days. GP breeders and non-breeders were housed as pairs of opposite or same sex, respectively. For this species, the time to first litters was 4.1±0.8 month and the pregnancies lasted about 68 days.

Female breeders gave birth to two litters each, with two exceptions. One NMR female was pregnant at least twice (ultrasonographically verified), but never gave birth to live offspring, and another gave birth to three litters, due to a pregnancy fathered by one of her sons. At time of sampling, NMRs and GPs reached an age of 3.4±0.5 and 0.9±0.1 years, respectively (Table S1).

**Table 1:**
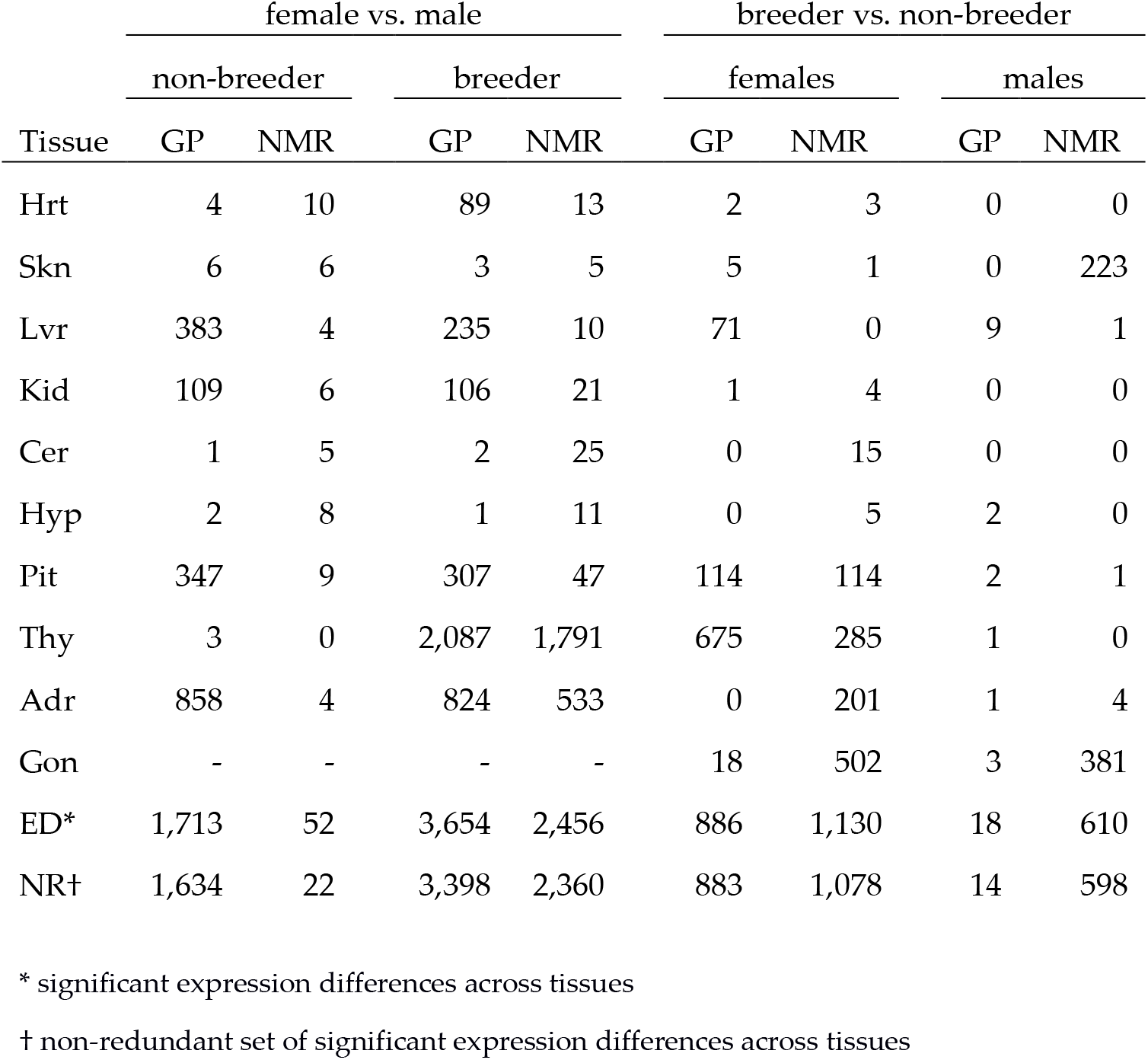
Numbers of DEGS identified in the different comparisons (FDR<0.01).

| Tissue | female vs. male |  |  |  | breeder vs. non-breeder |  |  |  |
| --- | --- | --- | --- | --- | --- | --- | --- | --- |
|  | non-breeder |  | breeder |  | females |  | males |  |
|  | GP | NMR | GP | NMR | GP | NMR | GP | NMR |
| Hrt | 4 | 10 | 89 | 13 | 2 | 3 | 0 | 0 |
| Skn | 6 | 6 | 3 | 5 | 5 | 1 | 0 | 223 |
| Lvr | 383 | 4 | 235 | 10 | 71 | 0 | 9 | 1 |
| Kid | 109 | 6 | 106 | 21 | 1 | 4 | 0 | 0 |
| Cer | 1 | 5 | 2 | 25 | 0 | 15 | 0 | 0 |
| Hyp | 2 | 8 | 1 | 11 | 0 | 5 | 2 | 0 |
| Pit | 347 | 9 | 307 | 47 | 114 | 114 | 2 | 1 |
| Thy | 3 | 0 | 2,087 | 1,791 | 675 | 285 | 1 | 0 |
| Adr | 858 | 4 | 824 | 533 | 0 | 201 | 1 | 4 |
| Gon | - | - | - | - | 18 | 502 | 3 | 381 |
| ED* | 1,713 | 52 | 3,654 | 2,456 | 886 | 1,130 | 18 | 610 |
| NR† | 1,634 | 22 | 3,398 | 2,360 | 883 | 1,078 | 14 | 598 |

\* significant expression differences across tissues
† non-redundant set of significant expression differences across tissues

### Tissue and species are the major determinants of transcriptomes

To compare gene expression between reproductive statuses (breeder vs. non-breeder) in NMR and GP, we performed RNA-seq of ten different tissue samples (heart - Hrt, skin - Skn, liver - Lvr, kidney - Kid, cerebellum - Cer, hypothalamus - Hyp, pituitary - Pit, thyroid - Thy, adrenal - Adr, and gonads – Gon, represented by either ovary – Ova or testis – Tes) from 24 animals for each species (six males, six females per status; Fig. S1). Seven of the 480 samples (1.5%) had to be excluded for different reasons (Tables S2, S3). On average (±SD), we obtained per sample 27.6±3.6 million high-quality reads with 84.1±16.1% unique mapping rate (Table S4). The grand mean of pairwise Pearson correlation within the 40 replicate groups (2 statuses x 2 sexes x 10 tissues per sex) was 0.981±0.013 and 0.984±0.01 for NMR and GP, respectively, indicating high consistency between replicate samples (Table S5).

Based on these data, unsupervised hierarchical clustering gave a similar cluster hierarchy of tissues for both species (Fig. S2). Brain tissues are grouped (Pit as a sister group to Cer and Hyp); Kid and Thy are sister groups to the cluster of Adr and Ova. The results are confirmed by principle component analysis, separating tissues by the first and species by the second component (Figure 1). At this level of analysis, ovary was the only tissue, which showed a separation of samples with respect to breeding status. Together, this indicates that: (i) tissue source is dominant over other biological variables such as species, sex and status and (ii) the impact of sex and status on transcriptome profiles is subtle.

**Figure 1:**
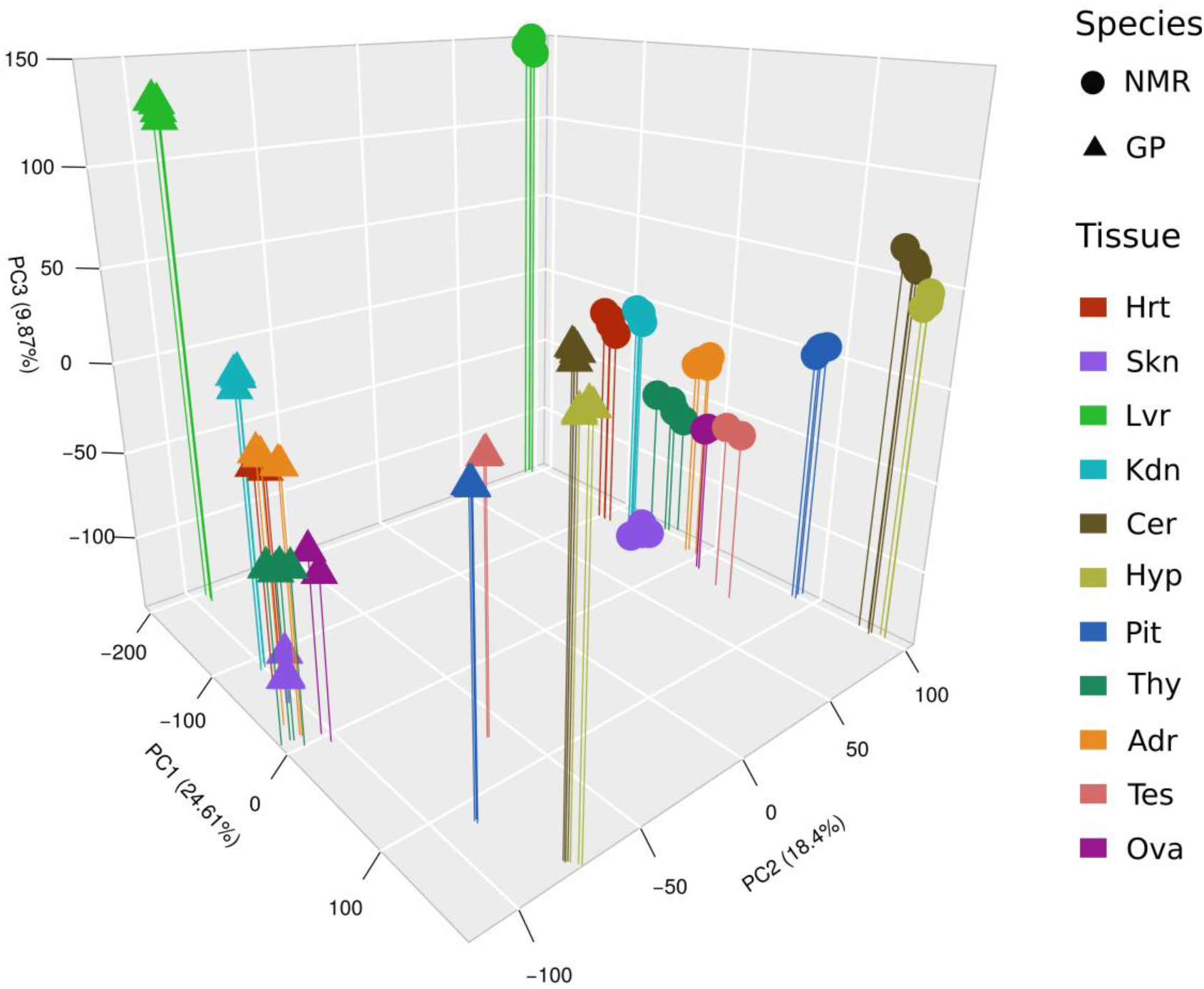
Principle component (PC) analysis of groups based on mean expression levels (four groups per tissue and species: 2 sexes x 2 statuses, except gonads). Tissues are separated by PC1 and PC3, species by PC2.

### Cross-species DEGs are enriched in aging-related genes

To further characterize the species differences between the long-lived NMR and the shorter-lived GP, we determined gene expression differences based on orthologous transcribed regions that show high sequence similarity. This filtering method avoided potentially misleading signals that may arise from assembly artifacts or the comparison of different transcript isoforms and identified 10,127 genes suitable for further analyses.

Across all tissues, we identified 18,000 significant expression differences (EDs; 9,651/8,349 higher/lower expressed in NMR) in 5,951 out of 10,127 genes (FDR<0.01, |log2FC|>2; Table S6, Supplemental Data S1-S11). Among genes that are consistently differentially expressed across all tissues, we identified aging-related candidates. E.g. *RRAGB* (Ras related GTP binding B, Figure 2A) and *TMEM8C* (transmembrane protein 8C) are higher expressed in NMRs. *RRAGB* interacts with mTORC1 complex [26, 27]. *TMEM8C* is essential for muscle regeneration [28] and might be linked to the resistance to muscle loss in aged NMRs [29, 30].

**Figure 2:**
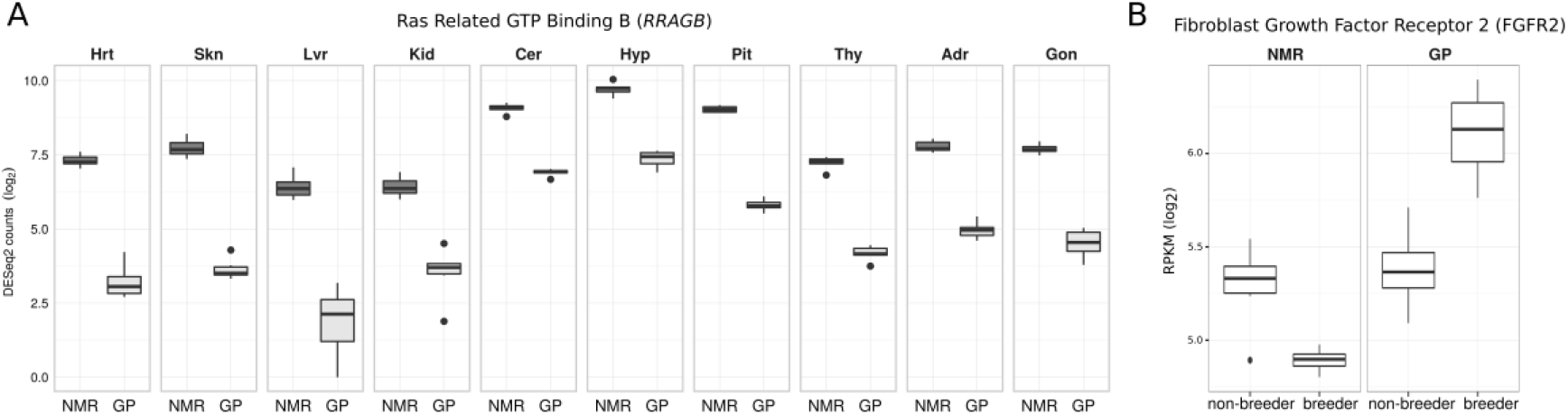
**(A)** Ras Related GTP Binding B (RRAGB) is consistently differentially expressed between species across all tissues. RRAGB is known to interact with mTORC1 complex [26, 27]. **(B)** Fibroblast Growth Factor Receptor 2 (FGFR2) shows opposing direction of expression between NMR and GP in breeders vs. non-breeders.

To further assess the association of cross-species DEGs with aging, we examined their overlap with aging-related genes of human and mouse obtained from the Digital Ageing Atlas (DAA) [31]. This test revealed a significant overlap with DAA containing 1,056 genes (17.74% of DEGs; p=0.006, Fisher’s exact test (FET); Table S7). The enrichment analysis of shared aging-related genes (Table S8) reveals that the top ranked GO term set is associated with lipid biosynthetic process (GO:0008610) (Fig. S3), a finding referring to existing links between lipid metabolism and lifespan [32, 33].

### Sexual differentiation and maturation in NMR are delayed until transition from worker to breeder

The DEGs between sexes were determined within the groups of non-breeders and breeders for each tissue and species (Table 1, Table S9, Supplemental Data S12-S47). GP non-breeder females vs. males (GP-N-FvM) show over all tissues except gonads 1,713 significant expression differences (EDs) in 1,634 genes (FDR<0.01, Table 1), primarily in Adr (858 DEGs), Lvr (383), Thy (347) and Kid (109). Between female and male GP breeders (GP-B-FvM) 3,654/3,398 EDs/DEGs were observed. These transcriptome data confirm a clear sexual differentiation among sexually mature GPs that further increases after onset of breeding. Breeders have 790 DEGs in common with non-breeders (p<2.2×10^−16^, FET; Figure 3A). Functional enrichment analysis of these shared genes reveals among the highest ranked GO sets immune system related terms (Fig. S5, Table S10).

**Figure 3:**
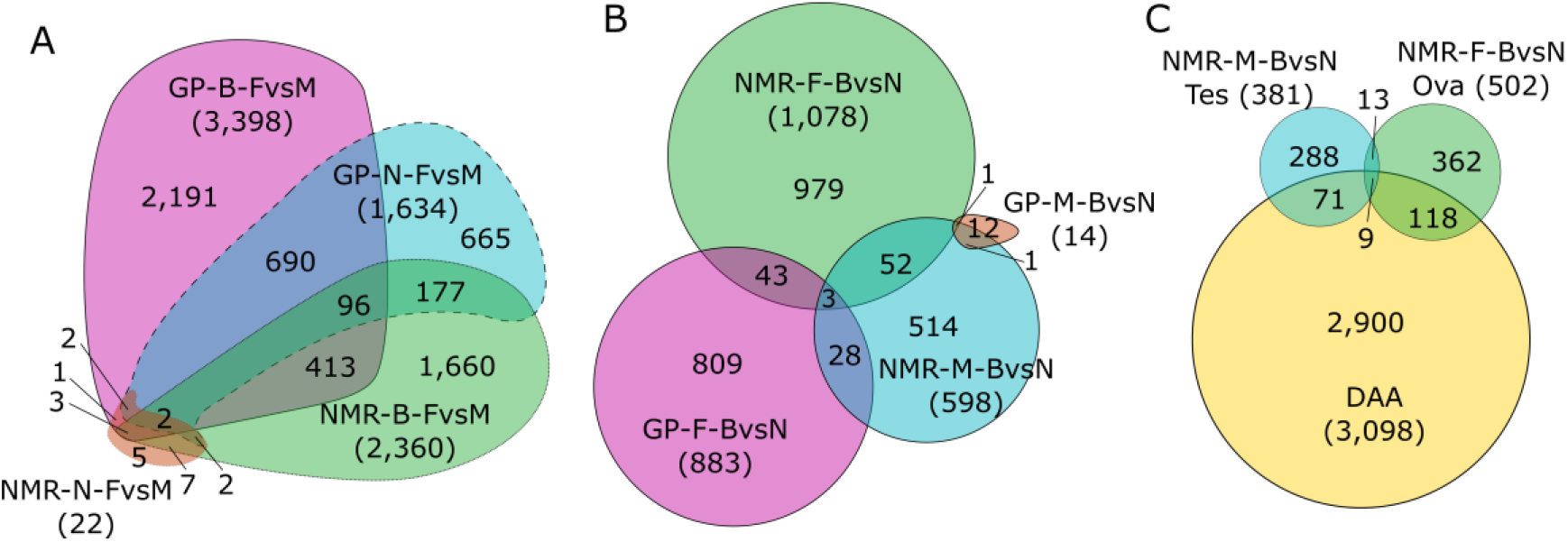
Euler diagrams showing overlaps of DEGs. **(A)** Female vs. male. **(B)** Breeder vs. non-breeder. **(C)** Gonads of NMR breeder vs. non-breeder and aging-related genes from the Digital Aging Atlas (DAA).

Similarly to GP-B-FvM, NMR-B-FvM showed 2,456/2,360 sex-related EDs/DEGs (FDR<0.01), mostly in Thy (1,791) and Adr (533). The overlap with GP-B-FvM is with 514 DEGs considerable but does not reach significance (p=0.062, FET; Figure 3A). Nevertheless, these data indicate basic similarities in sexual differentiation among breeders of both species.

Surprisingly, only 22 NMR-N-FvM EDs/DEGs were detected across all tissues (Table S11), indicating an only minor sex differentiation between NMR non-breeding females and males on the transcriptional level, consistent with the almost absent sexual dimorphism among non-breeding NMRs [23, 24].

### Status change of NMRs is accompanied by major changes in the endocrine system

The DEGs between breeders and non-breeders were determined within the same sex for each species (Table 1; Table S12, Supplemental Data S48-S87). Females showed a similar amount of EDs/DEGs in both species (GP-F-BvN: 886/883, NMR-F-BvN: 1,130/1,078) but have only 46 DEGs in common (Figure 3B). This is less than expected by chance, although not reaching significance (p=0.075, FET for depletion) and indicates that the molecular signature of the transition from female non-breeder to breeder is different in both species. E.g. in GP-F-BvN, only 18 DEGs are observed in Ova and none in Adr while in NMR-F-BvN, these tissues show most of the differences with 502 and 201 DEGs, respectively. Functional enrichment analysis of DEGs in NMR Ova identifies reproductive structure development (GO:0048608) as the highest ranked category (Fig. S6, Table S13). The same analysis in Adr revealed an obvious directionality in expression changes. DEGs are preferentially upregulated among highest ranked GO term sets (Fig. S7, Table S14), e.g. in reproduction (GO:0000003, 24 of 28) and endocrine system development (GO:0035270, 17 of 18). Tallying with this, Cer DEGs are enriched and upregulated in steroid metabolic process (GO:0008202, 6 of 6 upregulated) and response to hormones (GO:0009725, 7 of 7) (Fig. S8, Table S15).

In male GPs, status-related differences (GP-M-BvN) were almost absent across all tissues (only 18 EDs/14 DEGs). In contrast, NMR-M-BvN showed 610/598 EDs/DEGs, predominantly in Tes (381) and Skn (223). NMRs share 55 status-related DEGs in both sexes (p=0.008, FET; Table S16), while the few status-related changes in male GPs showed no overlap with those in females (Figure 3B). Among shared DEGs in NMRs, 10 genes involved in endocrine signaling were identified, including *SSTR3* (somatostatin receptor), *TAC4* (tachykinin), *PRDX1* (peroxiredoxin 1) and ACPP (acid phosphatase, prostate), as well as in general signaling via cAMP signaling (three genes) and through G-protein coupled receptors (four genes) further underlining that social status transition in NMR is associated with changes in the endocrine system.

### Mitochondrial genes show opposed expression changes in Tes and Skn after status change of NMR males

In NMR-M-BvN Tes, functional enrichment analysis revealed as highest ranked metabolism- and energy-related GO sets. DEGs enriched therein are mostly upregulated, e.g. lipid biosynthetic process (GO:0008610, 75 of 82 genes) and oxidation-reduction process (GO:0055114, 64 of 64) (Fig. S9, Table S17). Furthermore, we observed an upregulation of response to stimulus (GO:0050896, 79 of 106), in line with an upregulation of steroid metabolic process (GO:0008202, 28 of 28) included in lipid biosynthetic process set. In accordance with the dominance of energy-related processes, DEGs are enriched and preferentially upregulated in the top GO cellular component terms mitochondria (GO:0005739, 61 of 62) and peroxisomes (GO:0005777, 13 of 13) (Table S18). Together, this indicates increased demands of energy, e.g. to produce steroid hormones in Tes of NMR breeders.

Similar to Tes, Skn showed enrichment of energy-related processes (Table S19, Fig. S10). However, GO term sets in Skn are mostly downregulated, including energy derivation by organic compounds (GO:0015980, 36 of 38 genes) and oxidation-reduction process (GO:0055114, 42 of 46). Consistently, this includes genes associated with mitochondria (GO:0044429, 56 of 57) and respiratory chain (GO:0070469, 11 of 11).

The overlap between mitochondrial DEGs in Tes and Skn comprises 6 genes (p=7.27×10^−8^, FET; Figure 4). Among common genes *PINK1* (PTEN induced putative kinase 1) is 1.5-fold upregulated in Tes and 2.5-fold downregulated in Skn, indicating a role in regulation of mitophagy [34] in both tissues.

**Figure 4:**
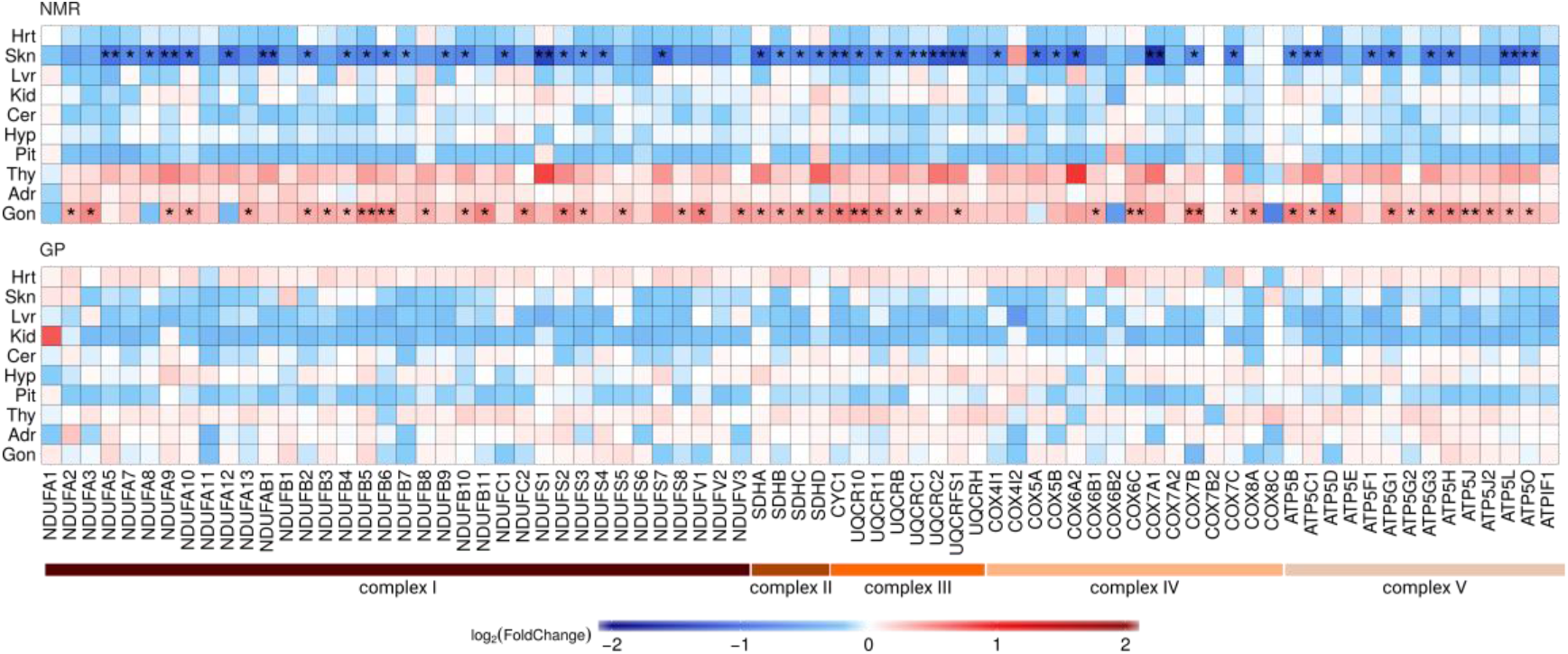
Expression changes of nuclear genes encoding for mitochondrial respiratory chain complexes in status change of male NMRs (top) and GPs (bottom). DEGs are indicated by asterisk (*: FDR<0.05, **: FDR<0.01). Only NMRs show significant expression differences (skn: 46 genes, mean fold-change 0.76; tes 46 genes, mean fold-change 1.89).

To follow up the mitochondria-related findings, the ‘mitonuclear transcript ratio’ was determined as the read count ratio of mitochondrial-encoded genes versus nuclear-encoded genes. It differs largely between tissues and species (Table S20). Hrt showed the highest mitonuclear ratio, with a minor difference between species (NMR 30.1%, GP 30.4%). Tes showed the lowest ratio, particularly in GPs (NMR 5.5%, GP 0.9%), and a 43.1% increase in NMR-M-BvN (Fig. S11). This increase is accompanied with an upregulation of nuclear genes encoding for mitochondrial respiratory chain complexes (Figure 4; Table S21). The expected increase in ROS is compensated by an on average 1.59-fold upregulation of eight antioxidant DEGs (Table S22). Consistently with functional enrichment analysis mentioned above, an opposing effect in Skn of NMR male breeders was observed, which showed a decline in mitonuclear ratio together with a downregulation of nuclear genes of the respiratory complexes (Figure 4). In line with the downregulation of the oxidative phosphorylation pathway (OXPHOS), a downregulation of antioxidant enzymes SOD2 (superoxide dismutase 2, 2.64-fold) and PRDX3 (peroxiredoxin 3, 2.14-fold) was observed. In general, downregulation of OXPHOS is linked to an extended lifespan [32, 35–37].

### NMR status-related DEGs are enriched in aging-related genes

Guided by the exceptional long life- and healthspan of NMR breeders, we searched our transcriptome data for molecular signatures relating to this phenomenon. First, we found that only NMRs show status-related DEGs that are significantly enriched for aging-related genes from DAA (Table S23): males in Skn (55 genes; q=0.0012, FET) and Tes (80 genes, q=0.01), and females in Ova (127 genes; q=1.2×10^−7^), Thy (59 genes, q=0.033) and Adr (43 genes, q=0.038). The significant overlap of 22 DEGs between NMR-F-BvN and NMR-M-BvN in Gon (p=0.0035, FET) contains nine aging-related genes (p=0.004, Figure 3C). In GP, only the non-redundant set of DEGs in GP-F-BvN showed a tendency of enrichment (160 genes, q=0.051), in contrast to NMRs, which showed enrichment in males (134 genes, q=4.5×10^−5^) and females (245 genes, q=7×10^−9^).

Second, we hypothesized that reproduction impacts the life expectancy of NMR and GP differently. Therefore, we searched for status-related DEGs that are shared in both species, but show opposing direction of expression. These genes might mark different coping mechanisms with the metabolic load of reproduction. As described above, the overlap of DEGs between species is very low (Table S24). Nevertheless, opposing direction of expression change can be observed in Ova (1 of 2 shared DEGs), and female thyroid (8/8) and testis (1/1). E.g. the fibroblast growth factor receptor 2 gene (*FGFR2*), linked to aging (AgeFactDB) [38], is in Ova downregulated in NMRs, but upregulated in GPs (Figure 2B).

Third, and based on the assumption that status-related EDs with the greatest interspecies difference (regardless of directionality) have an impact also on species-specific aging trajectories of breeders, we determined enrichment of aging-related genes in the upper quintile of those genes (Table S25). In males, we found significant enrichments of DAA genes in Skn (q=7.04×10^−7^) and Tes (q=0.0025); in females in Skn (q=0.0064), Hrt (q=0.0024), Pit (q=0.0053) and Ova (q=0.0013). Further functional enrichment analysis of these aging-related gene sets reveals differences between sexes. In males, the non-redundant set of genes shows enrichment for lipid metabolism (GO:0006629), energy metabolism (energy derivation by oxidation of organic compounds, GO:0015980; mitochondrial ATP synthesis coupled proton transport, GO:0042776), glutathione metabolic process (GO:0006749) and immune system (innate immune response-activating signal transduction, GO:0002758) (Table S26, Fig. S12). Females showed enrichment in positive regulation of tumor necrosis factor production (GO:0032760) and negative regulation of programmed cell death (GO:0043069) (Fig. S13, Table S27).

### Status related changes in NMR contradict the disposable some theory

Moreover, we assessed the connection of cross-species DEGs with expression changes that are associated with status change in each species. Based on DEGs shared in two comparisons (NMR vs. GP and breeder vs. non-breeder, FDR<0.05), we performed a correlation analyses of fold changes in each species. We hypothesized – in agreement with the disposable soma theory of aging – a negative impact of reproduction on lifespan for GP and – in contradiction 15 to this theory – an inverse effect for NMR. This was confirmed by opposing correlations (combined p=4.2×10^−9^ (Lancaster procedure [39]), negative correlation for GP and positive correlation for NMR (Figure 5). This means that DEGs with higher expression in NMR than GP are preferentially upregulated in NMR breeders compared to non-breeders, and vice versa; in other words, a transcriptome pattern associated with longevity (NMR vs. GP) is reinforced by the status transition (breeder vs. non-breeder) and thus may contribute to the exceptional long life- and healthspan of NMR breeders.

**Figure 5:**
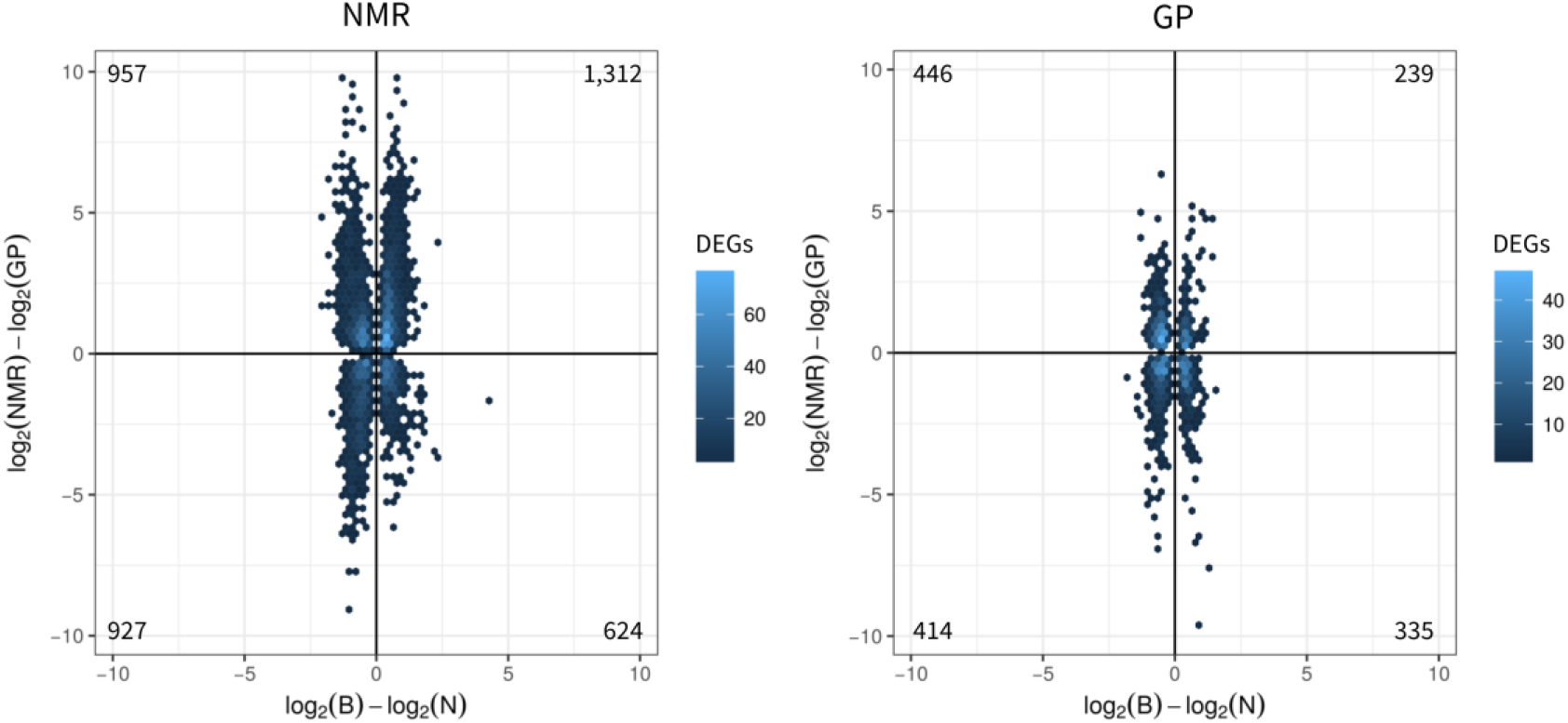
DEGs (FDR<0.05) occurring in both comparisons: cross-species (NMR vs. GP, y-axes) and status change GPs (breeder vs. non-breeder, x-axes) separately for (A) NMRs and (B). Correlation analysis between species shows opposing correlation (Lancaster procedure [39], p=4.2×10^−9^), while status-related DEGs in NMR are positively correlated with cross-species DEGs (DEGs=3,820; spearman correlation=0.17, p=3.2×10^−27^), status-related DEGs in GP show a negative correlation (1,434; −0.1; 1.8×10^−4^).

## Discussion

In our comparative study of breeders vs. non-breeders of the eusocial long-lived NMR and the polygynous and shorter-lived GP, we accumulated a comprehensive set of transcriptome data to gain insights into molecular networks underlying naturally evolved interspecies differences in sexual maturation and links between reproduction and aging. Both species are able to breed year-round and produce four to five litters per year [40, 41]. Both species have a similar average gestation period of 70 days, which is long compared to similarly sized species. Notably, NMRs produce on average 10.5 offspring per litter, which is twice the number of offspring produced by similarly sized rodents and more than three times higher compared to GPs (average 3.2 offspring) [40, 41]. This underscores the apparent contradiction of the NMR queen’s enormous metabolic load and extraordinary long life- and healthspan [4] to the disposable soma theory of aging [15], indicating that natural ways to extended healthspan remains to be uncovered in NMRs.

A first study to identify adaptations to unique NMR traits at the transcriptome level compared liver gene expression of young adult non-breeding male NMRs and mice [42]. Higher NMR transcript levels were observed for genes associated with oxidoreduction and mitochondria. This present study, more comprehensive in several aspects (sex, breeding status, numbers of animals and tissue samples), is based on a comparison of NMR vs. GP, which are phylogenetically closer than NMR and mouse. It revealed that between NMR and GP 58.8% of the analyzed genes are differentially expressed and that these DEGs are significantly enriched for aging-related genes. Among the latter, the main functional commonality is their association with lipid metabolism. Links of this molecular network to longevity of NMR were also obtained by a parallel proteome comparison of NMR and GP livers [43]. This study also shows that NMR liver mitochondria exhibit an increased capacity to utilize fatty acids.

In respect to sex-specific molecular signatures among either breeders or non-breeders in NMR and GP, the main finding is a nearly complete absence of significant transcriptional differences between sexes in non-breeding NMRs. This fits their grossly identical morphology and identical behavior in stable colonies [25]. This is in stark contrast to non-breeding GPs of an even younger age where we observed more than a thousand DEGs. GP non-breeders and breeders share a large and highly significant number of sex-related DEGs. Among others, these DEGs are enriched in GO terms related to steroid metabolism and immune system. The effect of gonadal steroids on the immune system is well described in GPs and other mammals [44]. After separation of NMR non-breeders from their colony, sexes became not only distinguishable by morphology and behaviour [22, 25], but also by gene expression. This differentiation on transcriptional level provides further molecular support for the previously described suppression of sexual maturation in non-breeding adult NMRs by social stress [8, 22] and identified major changes in the endocrine system after status change in NMRs but not in GPs (Supplement Text S1). Notably, we found no significant differences in gonadotropin-related genes (Supplement Text S2), indicating similar transcript turnovers in both non-breeders and breeders, and in line with previous results indicating that LH is stored in non-breeders, ready to be released upon GnRH signalling [45].

Glucocorticoids have been linked to stress, reproduction and social behavior in a variety of species, including members of muroidae, primates and cichlids [46–48]. In NMRs, however, correlation between social status and urinary cortisol is not clear and seems to depend on colony stability [49, 50]. Here, we observed a significant upregulation of *NR3C1* (glucocorticoid receptor) in Tes of NMR male and Thy of female breeders (Supplemental Data S55/67). Interestingly, this is in line with elevated expression of glucocorticoid receptor in Tes in African cichlid breeders. Males can reversibly change between dominant and subordinate phenotypes [51]. Similar to NMRs, only dominant phenotypes are reproductively active. In male African cichlids, moreover, aggression is negatively correlated with expression of *SSTR3* (somatostatin receptor 3) in Tes. Similarly, we observed in NMR breeder Tes significant downregulation of *SSTR3* (Supplemental Data S67). This indicates that *SSTR3* may be associated with social dominance in NMRs as well.

As our study was primarily motivated by the exceptional long life- and healthspan of NMR breeders, we searched for evidence indicating that NMR status change has an impact on genes involved in aging. We found enrichment of aging-related genes in the non-redundant DEG sets of males and females, as well as enrichment in most tissues with at least 50 DEGs (male Skn and Tes; female Ova, Thy and Adr). This contrasts with our observations in GPs, which showed only a tendency of aging-relation for the non-redundant set of status-related DEGs in females.

Further, we observed significant tissue-specific changes in OXPHOS of male NMR breeders. While Tes showed an upregulation of nuclear-encoded mitochondrial genes and a respective increase in mitonuclear transcript ratio, Skn showed the opposite. Moreover, we observed significant enrichments of genes involved in fatty acid metabolism among status-related DEGs in both NMR tissues. Consistent with the role of mitochondria in lipid homeostasis and the observed directionality of changes in OXPHOS, fatty acid metabolism DEGs in Tes were preferentially up- and in Skn downregulated. While the increased mitochondrial activity in Tes probably complies with demands of energy for the production of sex steroids and their anabolic effect on physiology, such as growth of testis [17], the observed changes in Skn may indicate a link to the extraordinary healthspan of NMR breeders. Previously, it was observed that inhibition of complex I activity during adult life prolongs lifespan and rejuvenates the tailfin transcriptome in short-lived fish [36]. Enhanced lipid metabolism and reduced mitochondrial respiration were also linked to NMR longevity in a parallel liver proteome study comparing NMR vs. GP and old vs. young NMR [43].

Finally, we performed a correlation analysis between species (NMR vs. GP) and status (breeder vs. non-breeder) EDs confirming the basic hypothesis of the present work: in contrast to GP and in line with recent demographic studies performed in NMRs [14], the transition into breeders results in molecular signatures linked to extended life- and healthspan only in NMRs. Genes which are higher or lower expressed in NMR compared to GP are also preferentially up- or downregulated in NMR breeders (positive correlation) opposite to GPs (negative correlation). In other words, the positive correlation in NMR contradicts the disposable soma theory of aging, as EDs contributing to a long lifespan (higher/lower expression in NMR than GP) are preferentially increased in NMR breeders compared to non-breeders, while diminished in GPs as expected by this theory.

Taken together, our comparative transcriptome analysis of breeders versus non-breeders of the eusocial, long-lived NMR versus the polygynous and shorter-lived GP identifies molecular networks underlying socially regulated sexual maturation and naturally evolved extended life- and healthspan that encourage further functional and mechanistic investigations of these extraordinary NMR phenotypes.

## Materials & Methods

### Animals

*Naked mole-rats.* NMR colonies were kept inside a climatized box (2×1×1 m) in artificial burrow systems, consisting of eight cylindrical acrylic glass containers (diameter: 240 mm height: 285 or 205 mm). The latter functioned as variable nest boxes, food chambers or toilet areas, and were interconnected with acrylic tubes having an inner diameter of 60 mm. Husbandry conditions were stabile during the entire experimental period of 22 months. Temperature and humidity were adjusted to 27.0±2.0°C and 85.0±5.0%, respectively. In general, the NMR colonies were kept in darkness except for 2 to 4 hours of daily husbandry activities. Fresh vegetable food was provided daily and *ad libitum*. In addition, commercial rat pellets (Vita special, Vitakraft GmbH, Bremen, Germany) were fed as an additional source of protein and trace elements.

To turn them into breeders, randomly selected non-breeding animals derived from two long-term (>4 years) established colonies of more than 50 individuals were separated and paired with the opposite sex. As non-breeder controls, litter siblings of paired animals remained in their colonies as workers. After the lactation period of the second set of live offspring the tissue sampling was scheduled. To avoid further pregnancies in the females, male partners were removed and euthanized 8-10 days postpartum. The tissue collection in the females took place 40-50 days after the end of last pregnancy.

*Guinea pigs*. GPs (breed: Dunkin Hartley HsdDhl:DH, Harlan Laboratories, AN Venray, Netherlands) were housed in standardized GP cages (length: 850 mm, width: 470 mm, height: 450 mm) in breeding pairs plus offspring or in same-gender pairs of two. Commercial guinea pig pellets and commercial pet food hay (Hellweg Zooland GmbH, Berlin, Germany) were provided together with vitamin C enriched water *ad libitum*. Housing temperature and humidity were 18.0±2.0°C and 45.0±5.0%, respectively. A 12h light/dark regime was provided.

After an initial adaption period of 6 to 8 weeks the GPs were randomly divided in breeding pairs or in same-gender pairs of two. The offspring were separated from their parents after weaning (∼3 weeks postpartum). Tissue collection was scheduled after the lactation period of the second set of live offspring. To avoid further pregnancies in the females, male partners were removed between eleven days before and seven days after birth of the second litter. The tissue collection in the females took place 42-83 days after the end of last pregnancy.

For tissue collection, all animals were anaesthetized by 3% isoflurane inhalation anaesthesia (Isofluran CP, CP-Pharma, Burgdorf, Germany) and euthanized by surgical decapitation. Animal housing and tissue collection at the Leibniz Institute for Zoo and Wildlife Research was compliant with national and state legislation (breeding allowance #ZH 156; ethics approval G 0221/12 “Exploring long health span”, Landesamt für Gesundheit und Soziales, Berlin).

### Sample collection, RNA Isolation and Sequencing

For *de novo* transcriptome assembly, animals were euthanized and ten tissue samples (heart – Hrt (NMR only), skin - Skn, liver - Lvr, kidney - Kid, cerebellum - Cer, hypothalamus - Hyp, pituitary - Pit, thyroid - Thy, adrenal - Adr, and gonads - Gon (testis – Tes /ovaries - Ova)) were collected from NMR and GP individuals, as described previously [52]. Strand-specific RNA-seq were prepared using the TruSeq Stranded RNA LT Kit (Illumina), and 200-nt reads were obtained using a HiSeq2500 (Illumina), as described previously [52].

For expression analysis, the same ten tissues were collected from NMR and GP breeders and non-breeders. RNA was purified as described above. Library preparation was done using Illumina’s TruSeq RNA Library Prep Kit v2 kit following the manufacturer’s description. Quantification and quality check of the libraries was done using Agilent’s Bioanalyzer 2100 in combination with a DNA 7500 Kit (both Agilent Technologies). Sequencing was done on a HiSeq 2500 running the machine in 51 cycle, single-end, high-output mode by multiplexing seven samples per lane. Demultiplexing and extraction of read information in FastQ format was done using the tool bcl2astq v1.8.4 (provided by Illumina).

### Data Analysis

*De novo* transcriptome assembly and annotation for GP was performed as described in [52]. Briefly, overlapping paired-end reads were joined into single fragments and then assembled by Trinity [53]. Gene symbols were assigned to the assembled transcripts by similarity to human transcripts using FRAMA [52].

As a reference for RNA-seq data mapping the public NMR (Bioproject PRJNA72441) [54] and GP genomes (UCSC, cavpor3) were used. Reference transcript sets of NMR and GP were mapped to the corresponding genome in two steps: BLAT (v36) [55] was used to identify the locus and then SPLIGN (v1.39.8) [56] was applied to splice align the transcript sequence within BLAT locus. RNA-seq data were aligned to the corresponding reference genome utilizing STAR (v2.4.1d) [57] with a maximum mismatch of 6% and a minimum aligned length of 90%. Reads mapped to multiple loci were discarded. Gene expression was quantified using HTSEQ (v0.6.1p1) [58] based on the aligned reference transcripts. The pairwise Pearson correlation between biological replicates was calculated based on 16,339 and 16,009 genes in NMR and GP, respectively (Table S5).

The ‘mitonuclear transcript ratio’ was calculated as the read count ratio of 13 mitochondrial-encoded genes versus all nuclear-encoded genes.

PosiGene was applied to the transcriptome of human, NMR and GP with the parameter ‘-prank=0 -max_anchor_gaps_hard=100 -rs=NMR’ to determine orthologous transcribed regions in NMR and GP having a protein identity >70%. RNA-seq data were aligned to the corresponding transcriptomes utilizing bowtie2 (2.2.9) [59] with the parameter ‘-very-sensitive-local’.

DESeq2 (v1.6.3) [60] was used to identify DEGs. A false discovery rate <0.01 (FDR; Benjamini Hochberg corrected p values, [61]) was used for the identification of significant DEGs.

Gene Ontology analysis was performed using the web interface of GoMiner (Database build 2011-01) based on the functional annotation of human genes (UniProt) [62]. A FDR<0.05 was used for the identification of significant GO terms that were summarized by REVIGO (parameter SimRel=0.5) into non-redundant GO term sets [63]. GO term sets were then ranked by number of summarized GO terms and number of changed genes. KEGG analysis was performed using Fisher’s exact test and significant pathways were identified using FDR<0.05. KEGG results were redundant to Gene Ontology analysis and therefore not shown.

Overlap between gene sets was determined with Fisher’s exact test (FET) using the one-sided option. Generally, we tested for enrichment if not otherwise stated.

We obtained 3,009 aging-related genes in human and mouse from the Digital Ageing Atlas (DAA) [31]. The corresponding counterparts in the NMR (2,588) and GP (2,539) were used for enrichment analysis and results were corrected for multiple testing (FDR). P-values corrected for multiple testing are indicated by q and nominal p-values by p.

To examine the connection between reproduction and aging in both species, we determined the difference in log2-fold-change (breeders vs. non-breeders) of NMR and GP. For fold-changes moving in opposite directions between species, we calculated the absolute difference 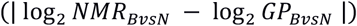, and for fold-changes moving in the same direction, higher fold-changes in NMR-BvN were rewarded 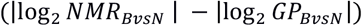. The 20%-quantile of genes having the greatest difference was determined separately for (i) the complete gene set and for genes showing (ii) opposing and (iii) unidirectional fold-changes. All sets were tested for enrichment of aging-related genes. Statistical analyses were performed in R (version v3.1.2).

## Data Availability

RNA-seq data for gene expression profiling were deposited at Gene Expression Omnibus (GSE98719). RNA-seq data for *de novo* assembly were deposited at Sequence Read Archive (SRP104222, SRP061363) and the corresponding gene collection is available as a gff3-file (ftp://genome.leibniz-fli.de/pub/nmr2017/).

## Acknowledgements

We like to acknowledge Angelika Kissmann and Jette Ziep for their professional help with the animal husbandry. We also thank Michaela Wetzel for her logistic help during tissue sampling and storage as well as for her support with the animal husbandry. We are grateful to Ivonne Görlich for excellent technical assistance as well as Debra Weih and Steve Hoffmann for critical reading of the manuscript.

## Funding

This research was supported by the Leibniz-Gemeinschaft through the Senatsausschusswettbewerb (SAW), grant SAW-2012-FLI-2, and the Deutsche Forschungsgemeinschaft (DFG), grant PL 173/8-1.

## Competing interests

None declared.

## Author Contributions

TBH, MP and KS conceived the project. SH, TBH, MB, MG and KS performed the animal study, sampling and sequencing experiments. Data analysis and interpretation were performed by MB, KS, SH, AS, HK, MP and TBH. The manuscript was written by MB and KS, and revised by all authors. The manuscript submitted by MB; all authors reviewed and approved the submitted manuscript. TBH and MP are joint senior authors.

## Abbreviations

DEG: differentially expressed gene
ED: expression differences
FDR: false discovery rate
FET: Fisher’s exact test
GO: Gene Ontology
GP: guinea pig
GP-B-FvM: comparison of GP Breeder Females vs. Males
GP-F-BvN: comparison of GP Female Breeders vs. Non-breeders
GP-M-BvN: comparison of GP Male Breeders vs. Non-breeders
GP-N-FvM: comparison of GP Non-breeder Females vs. Males
NMR: naked mole-rat
NMR-BvN: comparison of NMR Breeders vs. Non-breeders (both sexes)
NMR-B-FvM: comparison of NMR Breeder Females vs. Males
NMR-F-BvN: comparison of NMR Female Breeders vs. Non-breeders
NMR-M-BvN: comparison of NMR Male Breeders vs. Non-breeders
NMR-N-FvM: comparison of NMR Non-breeder Females vs. Males

## Supplementary Table Legends

**Table S1.** Age at death for NMRs and GPs.

**Table S2.** Number of analyzed biological replicates per group.

**Table S3.** Reason for exclusion of samples from further analysis.

**Table S4.** Number of uniquely aligned RNA-seq reads.

**Table S5.** Mean pairwise Pearson correlation coefficients between biological replicates in (A) naked mole-rat and (B) guinea pig.

**Table S6.** Number of differentially expressed genes (FDR < 0.01) between NMR and GP (A) without logFC threshold and (B) with logFC threshold (|logFC| > 2).

**Table S7.** DEGs in cross-species comparison that show an overlap with DAA.

**Table S8.** Gene set enrichment analysis for cross-species DEGs (FDR < 0.01, |logFC| > 2) that are age-related (DAA).

**Table S9.** Number of sex-related (female vs. male) differentially expressed genes in (A) non-breeder and (B) breeder (FDR < 0.01).

**Table S10.** Functional enrichment analysis of 790 DEGs shared between GP-B-FvM and GP-N-FvM.

**Table S11.** Gene description of sex-related DEGs in NMR-N-FvM.

**Table S12.** Number of status-related (breeder vs. non-breeder) differentially expressed genes in (A) females and (B) males (FDR < 0.01).

**Table S13.** Functional gene set enrichment analysis of DEGs in Ovr of NMR-F-BvsN.

**Table S14.** Functional gene set enrichment analysis of DEGs in Adr of NMR-F-BvsN.

**Table S15.** Functional gene set enrichment analysis of DEGs in Cer of NMR-F-BvsN.

**Table S16.** DEGs in intersection between NMR-F-BvsN and NMR-M-BvsN.

**Table S17.** Functional gene set enrichment analysis of DEGs in Tes of NMR-M-BvsN.

**Table S18.** Functional enrichment analysis in cellular components of DEGs in Tes NMR-M-BvsN.

**Table S19.** Functional gene set enrichment analysis of DEGs in Skn of NMR-M-BvsN.

**Table S20.** Proportion of mitochondrial transcriptonal output.

**Table S21.** Differentially expressed nuclear genes (FDR<0.05) associated with mitochondrial respiratory chain complexes in Tes and Skn of NMR-M-BvsN.

**Table S22.** Antioxidant enzymes differentially expressed in Tes and Skn of NMR-M-BvsN (FDR<0.05).

**Table S23.** Enrichment of age-related genes (Digital Ageing Atlas) in status-related DEGs. Only tissues having at least 50 DEGs were tested for enrichment.

**Table S24.** Shared status-related genes between NMR and GP.

**Table S25.** FDR values (Fisher’s exact test) for overlap between Digital Ageing Atlas and upper quintile of status-related expression differences having the greatest interspecies difference. Three groups were investigated: (i) opposing and (ii) same direction of expression in both species as well as (iii) independent of direction. Further, in each group direction in each species was tested separately.

**Table S26.** Functional enrichment analysis of the non-redundant set of aging-related 20%-quantiles that show the greatest interspecies difference in males (Skn, Tes).

**Table S27.** Functional enrichment analysis of the non-redundant set of aging-related 20%-quantiles that show the greatest interspecies difference in females (Hrt, Pit, Ovr).

## Supplementary Figure Legends

**Figure S1:** Collected tissues exemplified in schematic figure of NMR.

**Figure S2:** Hierarchical clustering of gene expression profiles. The clustering tree shows a clear separation of tissues in both species, but less pronounced differences between sex and breeding status.

**Figure S3:** Top 15 highest ranked GO sets based on enrichment analysis of cross-species DEGs between NMR and GP. GO sets are ranked by number of summarized GO terms (x-axes) and number of DEGs (alongside bar).

**Figure S4:** Top 15 highest ranked GO sets based on enrichment analysis of sex-related DEGs that are shared between GP breeder and non-breeder. GO sets are ranked by number of summarized GO terms (x-axes) and number of DEGs (alongside bar).

**Figure S5:** Top 15 highest ranked GO sets based on enrichment analysis of sex-related DEGs in NMR breeder. GO sets are ranked by number of summarized GO terms (x-axes) and number of DEGs (alongside bar, together with number of up- and downregulated genes).

**Figure S6:** Top 15 highest ranked GO sets based on enrichment analysis of status-related DEGs in ovary of NMR females. GO sets are ranked by number of summarized GO terms (x-axes) and number of DEGs (alongside bar, together with number of up- and downregulated genes).

**Figure S7:** Top 15 highest ranked GO sets based on enrichment analysis of status-related DEGs in adrenal gland of NMR females. GO sets are ranked by number of summarized GO terms (x-axes) and number of DEGs (alongside bar, together with number of up- and downregulated genes).

**Figure S8:** Top 15 highest ranked GO sets based on enrichment analysis of status-related DEGs in cerebellum of NMR females. GO sets are ranked by number of summarized GO terms (x-axes) and number of DEGs (alongside bar, together with number of up- and downregulated genes).

**Figure S9:** Top 15 highest ranked GO sets based on enrichment analysis of status-related DEGs in testis of NMR males. GO sets are ranked by number of summarized GO terms (x-axes) and number of DEGs (alongside bar, together with number of up- and downregulated genes).

**Figure S10:** Top 15 highest ranked GO sets based on enrichment analysis of status-related DEGs in skin of NMR males. GO sets are ranked by number of summarized GO terms (x-axes) and number of DEGs (alongside bar, together with number of up- and downregulated genes).

**Figure S11:** Mitonuclear ratios in non-breeders and breeders per sex, tissue and species. Boxplots shows median, 2nd/3rd quartiles, whiskers extend to 1.5 the interquartile range and dots values outside this range. P-values were calculated using a two-tailed t-test; *: p<0.05, **: p<0.01.

**Figure S12:** Top 15 highest ranked GO sets based on enrichment analysis of the non-redundant set of aging-related 20%-quantiles that show the greatest interspecies difference in males (Tes, Skn). GO sets are ranked by number of summarized GO terms (x-axes) and number of DEGs (alongside bar).

**Figure S13:** Top 15 highest ranked GO sets based on enrichment analysis of the non-redundant set of aging-related 20%-quantiles that show the greatest interspecies difference in females (Hrt, Pit, Ovr). GO sets are ranked by number of summarized GO terms (x-axes) and number of DEGs (alongside bar).

## Supplemental Text S1

In NMR-M-BvsN Tes, this is indicated by a significant upregulation of receptors for androgens (AR, 1.3-fold) and gonadotrophic hormones (LHCGR, 1.8-fold) as well as an increase in steroid production. Specifically, we observe an upregulation of genes involved in the transport of cholesterol into mitochondria (STAR, steroidogenic acute regulatory protein, 2.8-fold; TSPO, translocator protein, 1.5-fold; CAV1, caveolin 1, 1.4-fold), the rate limiting step of steroidogenesis (1), and conversion of cholesterol into pregnenolone (CYP11A1, cytochrome P450 family 11 subfamily A member 1, 1.8-fold), which is essential for all steroids. Genes encoding for enzymes that produce testosterone (HSD17B1, hydroxysteroid 17-beta dehydrogenase 1, 1.4-fold), the more potent metabolite dihydrotestosterone (SRD5A1, steroid 5 alpha-reductase 1, 1.3-fold) and cortisol (CYP51A1, cytochrome P450 family 51 subfamily A member 1, 2-fold) are upregulated as well (Supplement Figure 10). The upregulation of steroidogenesis is accompanied by a significant increase in the mitonuclear ratio and enrichment of nuclear DEGs associated with mitochondria, including OXPHOS. Consequently, the load of reactive oxygen species (ROS) is presumably higher in NMR Tes, but is compensated by increase in antioxidant enzymes. NMR-F-BvsN Ova shows more subtle changes in steroidogenesis by increased estradiol production as indicated by significant upregulation of CYP19A1 (aromatase, 2.5-fold) and increased sensitivity to androgens and progesterogens as indicated by upregulation of corresponding receptors (Ova, AR, 1,4-fold; Pit, PGR, 1.6-fold). We find, however, substantial expression changes in two genes involved in the regulation of follicle-stimulating hormone secretion (2), INHA (inhibin alpha subunit, 19.2-fold) and INHBA (inhibin beta A subunit, 4.4-fold), indicating that ovaries of non-breeding NMRs lack preovulatory follicles. Further, enrichment of steroid metabolic process, response to hormones and prostate gland growth indicates active feedback signalling also in females and an impact on developmental processes. Increase in signalling is further supported by significant enrichment of gene products located in secretory vesicle (secretory granule lumen, GO:0034774) and on cell surface (GO:0009986).

## Supplemental Text S2

Although well expressed, we found no significant differences for gonadotropin-related genes (GNRH1, FSHB, CGA, LHB). This indicates a similar intracellular transcript turnover in non-breeders and breeders, and is consistent with previous results indicating that LH is stored in non-breeders, ready to be released upon GnRH signalling (1). Notably, GnIH (NPVF), a negative regulator of reproductive function suggested to have a role in reproductive immaturity in NMRs due to its decreased protein expression in brain of NMR breeder (2), shows tendentially decreased mRNA expression in NMR breeders (males: 0.37, females 0.13). Also, previously elevated diencephalon mRNA levels in NMR were reported for AR (androgen receptor) in male breeders and for CYP19A1 (aromatase), ESR1 (estrogen receptor 1) and PGR (progesterone receptor) in female breeders (3). Our corresponding Hyp data do not support those increases (log2FC<0.1). Nevertheless, we identified significantly elevated transcript levels of these genes both in male (Tes: AR; Supplemental Data S67) and female breeders (Pit: PGR, Supplemental Data S54; Ova: AR, CYP19A1, ESR1, PGR, Supplemental Data S57) supporting a complex status-related function of these genes. Furthermore, steroid feedback and GnRH secretion are integrated by brain GABA and glutamate signalling in mammals and cichlids (4). Although, we do not observe equivalent significant differences in the NMR brain, the data show an upregulation of genes coding for three GABA receptor subunits (GABRB3, GABRG1, GABRP) and one glutamate receptor subunit (GRIK2) in Tes (Supplemental Data S67), as well as two of three differentially expressed GABA (GABRB2, GABRG3) and three glutamate receptor subunits (GRIK1, GRIK2, GRIK4) in Ova and female Pit of NMR breeders (Supplemental Data S54, S57). Together, this suggests increased neuronal plasticity and/or activity predominantly in gonads of NMRs after becoming breeders.

In the testis of immature chicken, expression of ADIPOR1/2 (adiponectin receptor 1/2) is significantly lower than in mature animals (5). It has been hypothesised that these genes are involved in supporting the higher metabolic activity related to spermatogenesis, testicular steroid hormone production, and transport of spermatozoa and testicular fluid. In line with these observations, our results show a significant upregulation of ADIPOR2 in Tes of NMR breeders (1.7-fold; Supplemental Data S67).

The renin-angiotensin system predominantly involved in cardiovascular control has also been associated with reproduction in mice and human (6). Signaling through receptors coded by AGTR1 (angiotensin II receptor type 1) in males, and AGTR1/2 in females has been associated with fertility and stimulation of reproduction. In accordance, we observed a significant upregulation of AGTR1 and AGTR2 in gonads of NMR male and female breeders, respectively (Supplemental Data S67,S57).

